# NAT/NCS2-hound: A Webserver for the detection and evolutionary classification of prokaryotic and eukaryotic nucleobase–cation symporters of the NAT/NCS2 family

**DOI:** 10.1101/332452

**Authors:** A Chaliotis, P Vlastaridis, C Ntountoumi, M Botou, V Yalelis, P Lazou, E Tatsaki, D Mossialos, S Frillingos, GD Amoutzias

## Abstract

Nucleobase transporters are important for supplying the cell with purines and/or pyrimidines, for controlling the intracellular pool of nucleotides and for obtaining exogenous nitrogen/carbon sources for the metabolism. Nucleobase transporters are also evaluated as potential targets for antimicrobial therapies, since several pathogenic microorganisms rely on purine/pyrimidine salvage from their hosts. The majority of known nucleobase transporters belong to the evolutionarily conserved and ubiquitous NAT/NCS2 protein family. Based on a large-scale phylogenetic analysis that we performed on thousands of prokaryotic proteomes, we have developed a webserver that can detect and distinguish this family of transporters from other homologous families that recognize different substrates. We can further categorize these transporters to certain evolutionary groups with distinct substrate preferences. The webserver scans whole proteomes and graphically displays which proteins are identified as NAT/NCS2, to which evolutionary groups and subgroups they belong to and which conserved motifs they have. For key subgroups and motifs, the server displays annotated information from published crystal-structures and mutational studies pointing to key functional amino acids that may help experts assess the transport capability of the target sequences. The server is 100% accurate in detecting NAT/NCS2 family members. We also used the server to analyze 9109 prokaryotic proteomes and identified Clostridia, Bacilli, β- and γ-Proteobacteria, Actinobacteria and Fusobacteria as the taxa with the largest number of NAT/NCS2 transporters per proteome. An analysis of 120 representative eukaryotic proteomes also demonstrates the server’s capability of correctly analyzing this major lineage, with plants emerging as the group with the highest number of NAT/NCS2 members per proteome.

## Introduction

The NAT/NCS2 (Nucleobase-Ascorbate Transporter / Nucleobase-Cation Symporter-2) protein family encompasses ion-gradient driven transporters of key metabolites or anti-metabolite analogs with diverse substrate preferences, ranging from purine or pyrimidine permeases in various organisms to Na^+^-dependent vitamin C transporters in human and other mammals [1] [2] [3] [4] [5] [6]. Their additional function as providers of nitrogen/carbon source may also affect energy production, replication and protein synthesis through the salvage pathways for nucleotide synthesis [7,8][9]. In addition to their important direct role on the central metabolism of the cell, these and other nucleobase transporters have attracted interest as potential targets of purine/pyrimidine-based antimicrobials that could either be selectively routed into target cells to act as anti-metabolites or selectively inhibit an essential nucleobase transporter of the target cell [10–14]

This protein family is one of the 18 known families of the APC superfamily [15] and represents a subset of APC families which conform to a distinct structural/mechanistic pattern. The NAT/NCS2 transporters consist of 14 transmembrane segments (TMs) divided in two inverted repeats (7+7) and arranged spatially into a core domain (TMs 1-4 and 8-11) and a gate domain (TMs 5-7 and 12-14) [16]. The core domain contains all major determinants of the substrate-binding site, whereas the gate domain contributes to alternating access by allowing conformational rearrangements and providing major gating elements. The proteins probably function as homodimers and may use an elevator-like mechanism to achieve alternating access [17,18]. Similar structural features are described for transporters of two other APC families, the Sulfate Permeases (SulP) [19] and the Anion Exchangers (AE) which includes the well studied chloride/bicarbonate exchanger (band 3) of human erythrocytes [20].

The NAT/NCS2 is split phylogenetically in two subfamilies. The first one, COG2233 or NAT, contains bacterial and fungal permeases for purines (xanthine, uric acid), bacterial permeases for pyrimidines (uracil, thymine), plantal and mammalian broad-specificity uracil/purine permeases (not present in human), and the mammalian L-ascorbate transporters SVCT1 and SVCT2. Insight on the transport mechanism of this subfamily has been provided by high-resolution crystal structures for two members, the uracil permease UraA of *E. coli* [16,18] and the xanthine/uric acid permease UapA of *Aspergillus nidulans* [17], coupled with extensive mutagenesis studies on UapA [21], the xanthine permease XanQ of *E. coli* [1,22] and few other homologs [23,24]. The other subfamily, COG2252 or AzgA-like [25], contains bacterial, fungal and plantal permeases for salvageable purines (adenine, guanine, hypoxanthine) which are less well studied with respect to structure-function relationships [7,26].

Despite their importance, membrane transporters in general and the NAT/NCS2 family in particular are not so extensively studied to date as other categories of proteins are, due to the inherent difficulties in experimentation and in accurate prediction of their function [27] [28]. Based on a large-scale evolutionary analysis that we performed in this study, we have i) identified in prokaryotes the major evolutionary groups and subgroups, with distinct substrate specificities, ii) identified key motifs for each phylogenetic group and subgroup that are related to substrate specificity, iii) developed a webserver that utilizes all the above information to detect and classify at proteome-scale NAT/NCS2 transporters and iv) analyzed with this webserver 9109 prokaryotic and 120 Eukaryotic proteomes so as to investigate which evolutionary lineages are rich in these transporters. We expect that this type of analyses and the accompanying computational tool, which are lacking in general for other families of transporters, will facilitate the experimental study of new homologs, provide a practical tool for assignment of homologs into functionally-relevant associated subgroups and also improve their annotation in the databases.

## Materials and Methods

### Development of HMMs and Meme motifs for the family, subfamilies and evolutionary clusters

All the annotated sequences of the 2A APC superfamily (organized in 18 families) were obtained from TCDB [15]. For each of the 18 families we generated protein alignments with Muscle and Seaview [29,30] that were manually edited and then used to build a hidden markov model for each one of them with HMMER [31].

Next, 4442 Bacterial AND 213 Archaeal Proteomes were downloaded from UNIPROT (January 2017) [32]. Their protein sequences were scanned with the above 18 HMMs and thus, 8291 proteins of the NAT/NCS2 family were identified and retained for further analysis. Afterwards, close homologs were removed with the Blastclust software, using as cutoff 70% protein identity over 70% of sequence length. Thus 1355 NAT/NCS2 sequences were retained after this step.

Subsequently, these sequences were fed to the MEME software [33] so as to identify 14 motifs of length 14-21 or 18-25 amino acids each. Manual inspection of sequences with a very low number of motifs resulted in rejection of 14 sequences. Thus 1341 sequences were retained. These 1341 sequences were scanned again with the 14 MEME motifs, by MAST [33]. Custom Perl scripts were developed to obtain the motif presence/absence for each sequence as a vector of 0 and 1 values, based on detection with MAST (see supplementary folder Custom_scipts). The above vectors were clustered in MATLAB with the Clustergram function (default parameters – commands found in supplementary folder Custom_scipts). This first round of clustering revealed two major evolutionary subfamilies, designated SF1 and SF2 (see supplementary figure S1). The sequences of each subfamily were fed to another round of MEME motif detection with the same parameters as in the first instance. Again, 14 MEME-motifs were made for each subfamily. These were used with the MAST software to identify MEME-Motif content for each subfamily, and again, vectors of motif presence/absence were generated for each subfamily (and their clusters were manually inspected; see supplementary figures S2 and S3). All Meme/Mast results and analyzed sequences are found in the supplementary folder “Meme_Mast_motifs”.

Afterwards, the protein sequences of each subfamily separately were aligned and manually edited with Muscle and Seaview [29,30]. Furthermore, in each subfamily, sequences with experimental evidence of substrate specificity were added (eukaryotic ones as well). Phylogenetic trees were generated with the BioNJ method using the Poisson model and 1000 bootstraps. The two generated phylogenetic trees (for each of the two distinct subfamilies – see supplementary figures S4 and S5) were annotated and visualized in Archaeopteryx and Treedyn [34,35]. Subfamily 1 was organized in six major and four very small clusters. Subfamily 2, that was more homogeneous than subfamily 1, was organized in many small clusters. Hidden Markov Models were thus constructed for the NAT/NCS2 family, its two subfamilies and for each of the 6 major clusters in subfamily 1. For several of the small clusters in subfamily 2 that contained sequences with known substrates we also generated extra HMMs. In addition, we generated 14 MEME motifs for each subfamily and each of the 6 clusters in subfamily 1. All edited sequence alignments are organized in supplementary folder “Sequence_Alignments_edited”.

### Development and Evaluation of the server

All the above HMMs and MEME motifs were incorporated in a webserver, named NAT/NCS2-hound, that may scan protein sequences in FASTA format, identify members of this family and further classify them in the various subfamilies and clusters. The webserver is based on the Jhipster Application Framework (http://www.jhipster.tech/) that utilizes Angular Javascript Framework for the front-end and the Java language and Spring Framework for the back-end. The server is freely available at http://bioinf.bio.uth.gr/nat-ncs2/

The server and instructions for local installation are found in supplementary folder “Server_for_local_installation”.

Functional information for the various amino acids was obtained from several mutational studies [1,21,24] and from the structural studies on UraA [16,18] and UapA [17].

We performed an evaluation analysis, in order to assess the effectiveness of the NAT/NCS2-hound server. TCDB annotated transporters of the 18 families of the APC superfamily were used as bait to obtain best blast hits against bacterial reference proteomes downloaded from Uniprot. The best blast hit of a bait sequence was designated as a member of the family that its annotated (from TCDB) bait sequence belonged to. These retrieved best blast hit sequences constituted the evaluation set. Any of these sequences that had been used to train the HMMs were removed from the evaluation set.

Thus, we retrieved/retained 7799 APC sequences, of which 975 belonged to the NAT/NCS2 family. These were scanned by our server for detection and evolutionary classification. The server demonstrated 100% accuracy (100% sensitivity and 100% specificity) in detecting NAT/NCS2 family members and can further categorize them to the various evolutionary subgroups, display conserved motifs and relevant functional information/annotation.

In order to assess the distribution of NAT/NCS2 family in major taxonomic lineages, 9109 prokaryotic proteomes (downloaded from NCBI at March 2018) and 120 Eukaryotic Reference Proteomes (downloaded from Uniprot at March 2018) were scanned by our server. The presence of a minimum number of seven MEME motifs was required as a cutoff, to filter out any sequence fragments. All results and sequence IDs are found supplementary excel file1.

## Results and Discussion

### The NAT/NCS2 family is organized in two major subfamilies

An analysis of 1341 proteins, based on the presence/absence of conserved MEME motifs within the NAT/NCS2 family clearly revealed the presence of two distinct and major subfamilies (see supplementary figure S1). Previous phylogenetic analyses also revealed the presence of these two major subfamilies [7], in accordance with the presence of two COGs, designated as COG2233 (Xanthine/Uracil permease) and COG2252 (AzgA-like). Subfamily 1 (COG2233) consisted of 748 sequences and subfamily 2 (COG2252, AzgA-like) consisted of 593 sequences. The members of Subfamily 1 display a greater degree of sequence divergence among them, whereas members of subfamily 2 constitute a more homogeneous set of sequences (Supplementary figures S2-S3).

### Distinct Phylogenetic clusters within Subfamily 1

Further phylogenetic analysis of the more diverse members of Subfamily 1 reveals the clear presence of six major clusters (Clusters 1-6) and four minor clusters (see Figure 1). The incorporation of functionally annotated sequences from all kingdoms of life further helped us understand the substrate specificity profile of each cluster, whenever relevant information for representative homologs was available. The largest and most diverse cluster (Cluster 1) contains sequences that have been annotated to transport Xanthine or Uric Acid or both. The second largest cluster (Cluster 2) contains sequences that are known to transport Uracil or Uracil and Thymine. The third largest cluster (Cluster 3) contains the YbbY gene from *E.coli*, a homolog that is not functionally annotated in the databases but recent evidence suggests that it transports adenine, guanine and hypoxanthine (Botou and Frillingos, unpublished data). The fourth cluster (Cluster 4) contains functionally characterized sequences from Eukaryotes, but also encompasses functionally unknown homologs from Archaea as well as a few Bacteria. All the other clusters do not contain any sequences of known function.

**Figure 1.**
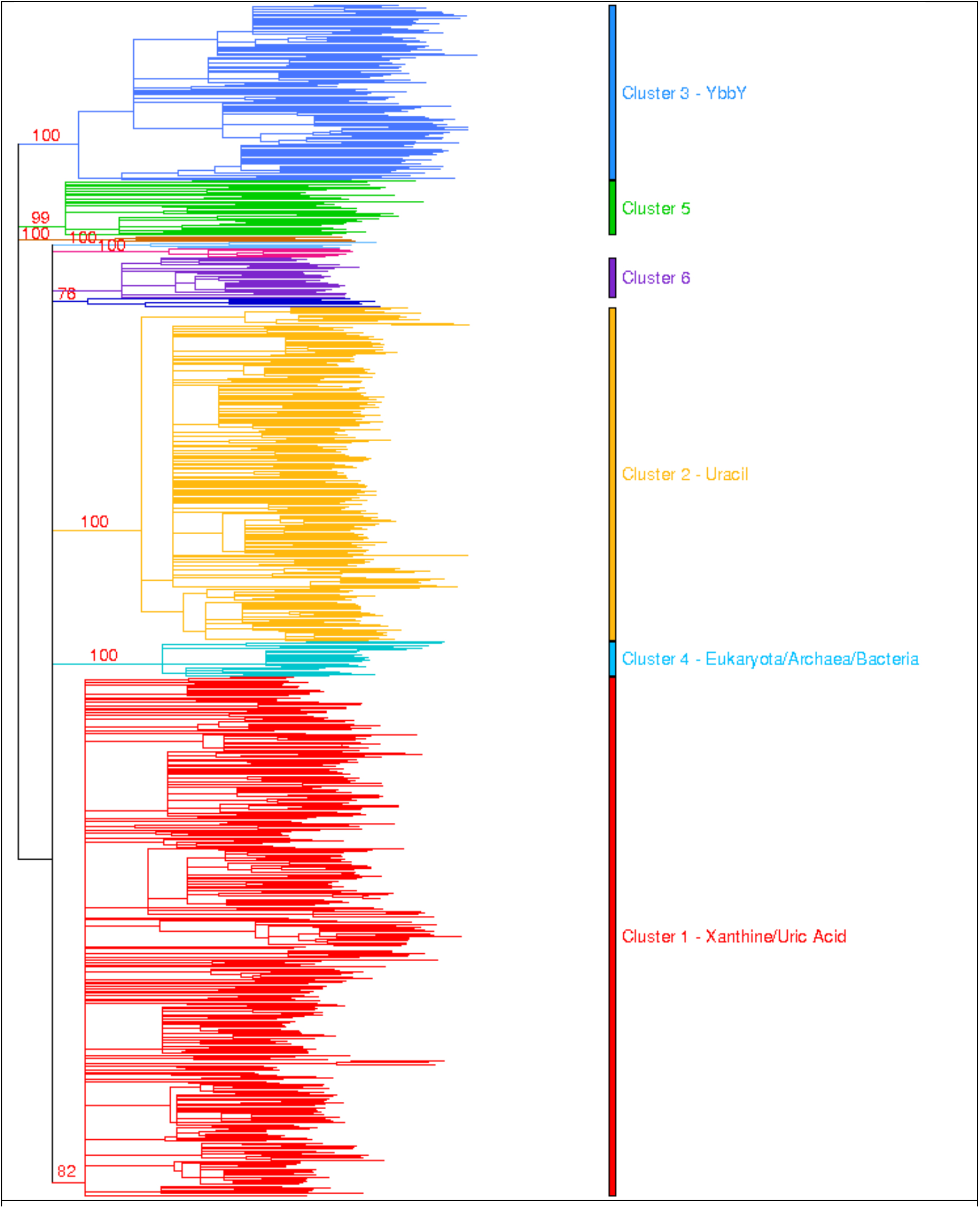
Phylogenetic tree of Subfamily 1 of the NAT/NCS2 family. The various phylogenetic clusters are depicted with different colors. Sequence redundancy was removed at a cutoff of 70% protein identity over 70% of sequence length.

Subfamily 2 is more homogeneous and is organized in many small clusters, with small differences among them. For several of those small clusters that contain sequences with known substrates we generated additional HMMs. A more detailed inspection of the various phylogenetic trees for each subfamily and each of the major 6 clusters within subfamily 1 are available in supplementary materials (Supplementary figures S4-S11).

### A web server for the detection and evolutionary classification of NAT/NCS2 family members

All the above evolutionary analyses, the HMMs and MEME motifs generated for the various subfamilies and evolutionary clusters have been used to develop a web server for the detection and evolutionary classification of NAT/NCS2 family members, named NAT/NCS2-hound. The webserver is freely available at http://bioinf.bio.uth.gr/nat-ncs2/. This webserver detects and distinguishes this family of transporters from the other 17 homologous families of the APC superfamily. Furthermore, it can categorize these transporters to certain subfamilies and clusters associated with distinct substrate specificities, based on the large-scale phylogenetic analysis that we performed on prokaryotic proteomes. For each one of them separately, the identified set of characteristic signature motifs is detected. Furthermore, for several key subgroups we have integrated information from published crystal-structures and mutational studies to help experts identify key functional amino acids and help them assess the transport capability of the scanned sequences. Nevertheless, this server does not function as a prediction tool of substrate specificity. The NAT/NCS2-hound server implements for this important family the same principles and computational protocol that were developed/implemented recently for another prokaryotic superfamily, the tRNA-synthetases [36].

The input for this server is a protein sequence or a proteome file in FASTA format. The webserver displays graphically (see figure 2) which proteins have been identified as NAT/NCS2, to which subfamily and cluster they belong to and which conserved motifs have been identified on the target proteins. For several key subgroups and motifs, the server further displays annotated (by our experts) information from published crystal-structures and mutational studies pointing to key functional amino acids of well-studied representative homologs.

**Figure 2.**
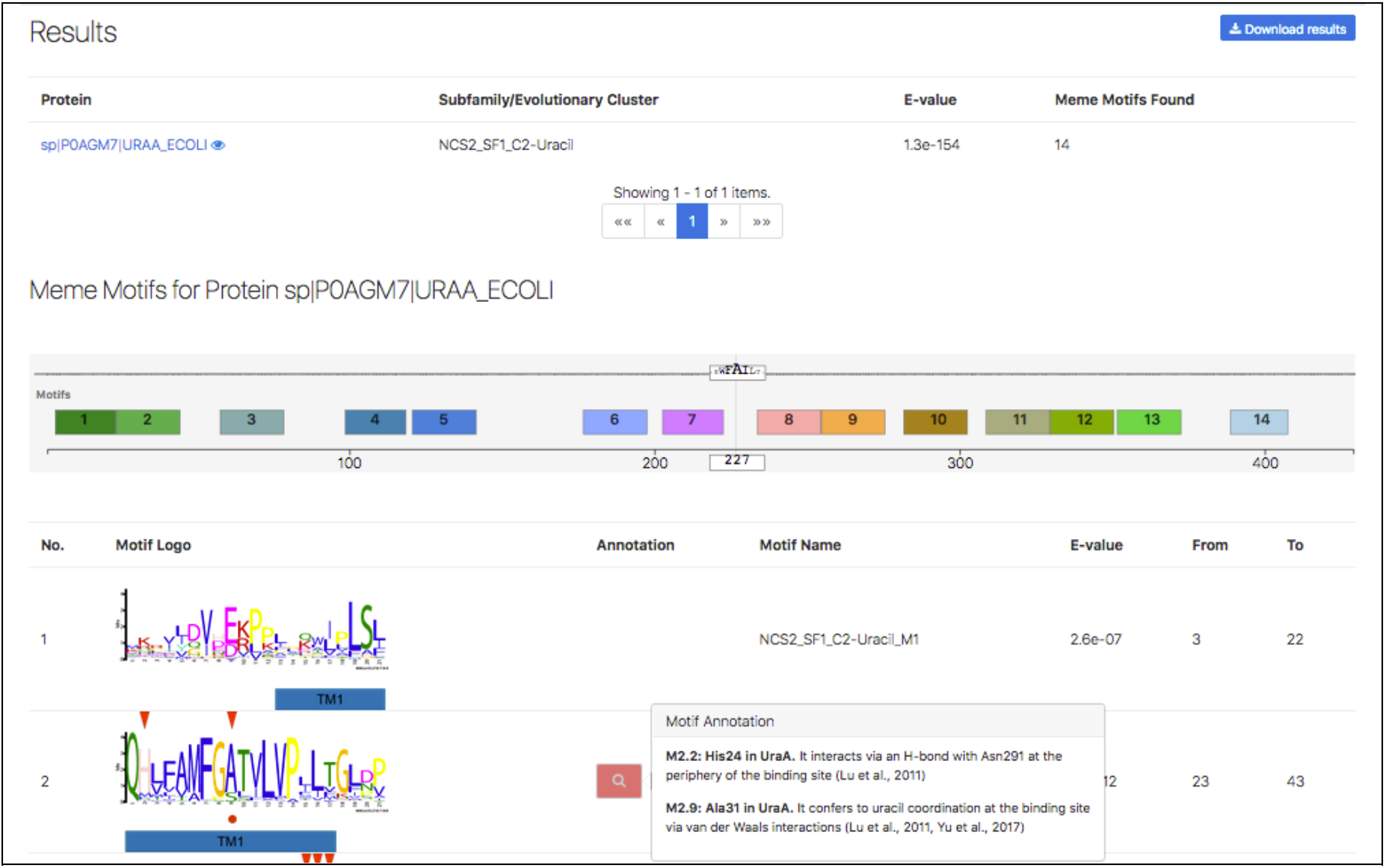
Display of results from the NAT/NCS2-hound server, including the best Hidden Markov Model that detects the protein sequence, the various MEME conserved motifs and any available functional information/annotation for specific sites in certain motifs.

The server has been evaluated against a dataset of 7800 homologous transporters of the APC superfamily, of which 975 belong to the NAT/NCS2 family and displayed 100% accuracy (100% sensitivity and 100% specificity).

### Distribution of the NAT/NCS2 family and subfamilies in Prokaryotes and Eukaryotes

In order to assess the distribution of NAT/NCS2 members in major taxonomic lineages, we scanned 9109 prokaryotic proteomes (from NCBI) and 120 representative Eukaryotic Proteomes (from Uniprot). As a filter, we only included in our analysis 29096 prokaryotic and 361 eukaryotic NAT/NCS2 proteins that had at least 7 Meme motifs each, to exclude small sequence fragments.

80% (7318/9109) of the prokaryotic proteomes had at least one NAT/NCS2 protein, based on our criteria, with 4 NAT/NCS2 proteins, on average. The lineages with the most proteins (on average) are Clostridia, Bacilli, β- and γ-Proteobacteria and Actinobacteria (see figure 3). The proteomes with the most transporters were *Clostridium bolteae* (with 14 members), several different γ-Proteobacteria (*Morganella morganii, Enterobacter lignolyticus, Citrobacter amalonaticus*) and *Bacillus megaterium* with 11 members each, followed by many other γ-Proteobacteria (such as *E.coli*) with 10 members each (see supplementary excel file 1; spreadsheet: Organisms). Subfamily 1 was more abundant than subfamily 2, comprising of 58% of all NAT/NCS2 proteins (paired t-test p-value =0). Also, Cluster 1 (Xanthine/Uric acid) and Cluster 2 (Uracil) from subfamily 1, comprised of 29% and 23% of the total number of NAT/NCS2 proteins detected, whereas Cluster 3 (YbbY) was the third largest one, comprising of 5% of the total number of NAT/NCS2 proteins.

**Figure 3.**
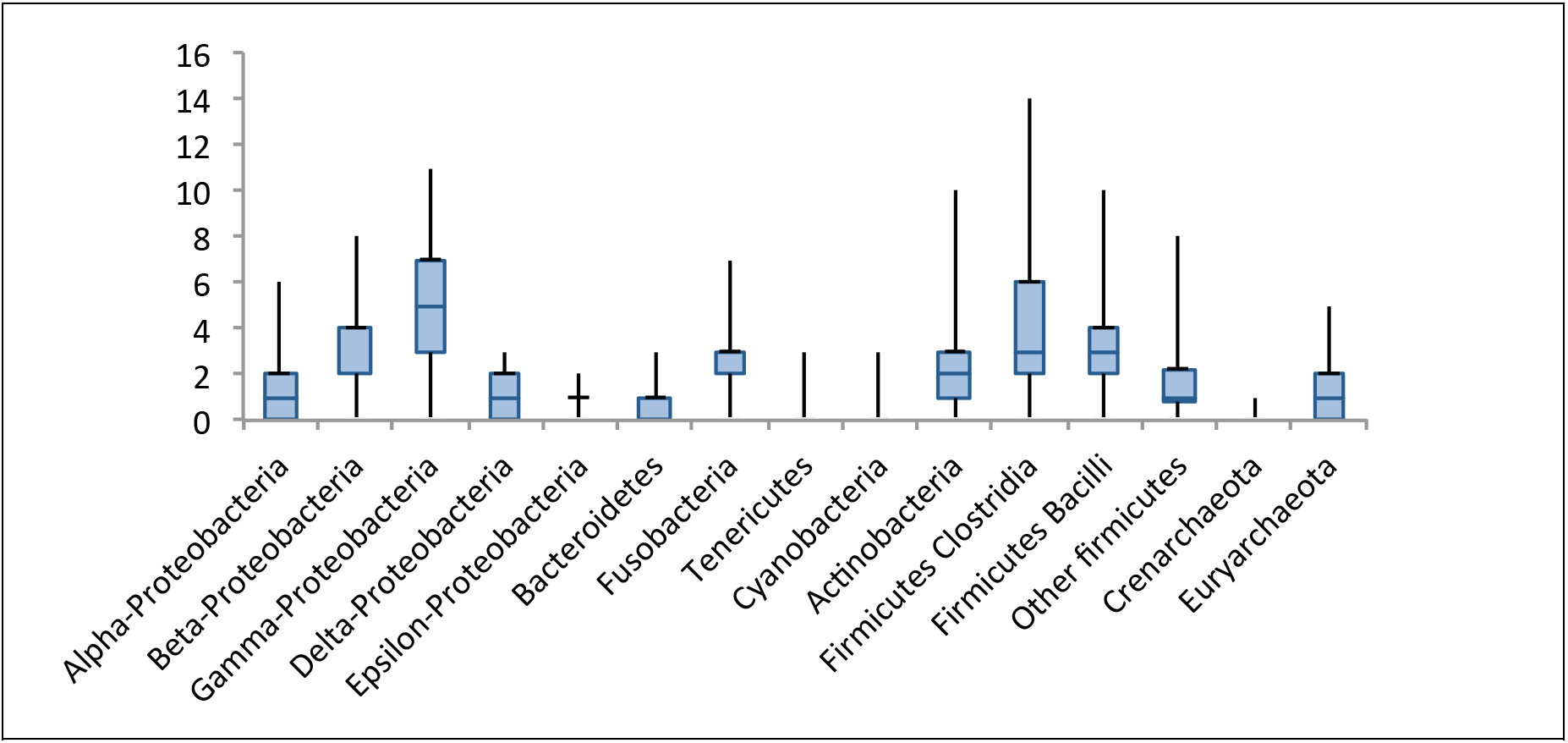
Boxplot of the distribution of NAT/NCS2 proteins per proteome, in major prokaryotic taxonomic lineages. The analysis was based on 9109 Bacterial and Archaeal proteomes. β and γ-Proteobacteria, Clostridia, Bacilli, Fusobacteria and Actinobacteria had the highest numbers of transporters per proteome (on average).

Furthermore, we did a survey on 61 proteomes from strains found in the human microbiome (https://www.hmpdacc.org/catalog/) and found that NAT/NCS2 homologs are enriched (3 per genome, on average) in bacteria of the gastrointestinal tract (see Supplementary excel file 2)

We also analyzed 120 representative proteomes from animals, fungi, plants and various unicellular Eukaryotes. 83% of the detected proteins were found to belong to subfamily 1 (the rest in subfamily 2), with the vast majority of them (66% of the total) in Cluster 4, followed by Cluster 1 (14% of the total) and Cluster 2 (3% of the total) (see supplementary excel file 3 for detailed results and analyzed sequences). As an additional validation step, our server detected and properly classified well-known and previously annotated NAT/NCS2 sequences in certain selected species, such as the human homologs SVCT1 and SVCT2 [5], the rat uracil/purine transporter rSNBT1 [4], the uracil/purine transporters AtNAT3 and AtNAT12 and adenine/guanine transporters AtAzg1 and AtAzg2 of *Arabidopsis thaliana* [3], and the xanthine/uric acid transporters UapA and UapC and adenine/guanine/hypoxanthine transporter AzgA of *Aspergillus nidulans* [2], Thus, although the development of the server was based on prokaryotic sequences, the server can successfully analyze eukaryotic sequences as well. This is attributed to the fact that all eukaryotic sequences are fully contained within evolutionary groups that are already present in Prokaryotes. Plants had the highest number of NAT/NCS2 members (10 per genome on average), followed by animals (3 on average), then by fungi (2 on average), whereas most (73%) of the unicellular eukaryotes had no NAT/NCS2 proteins. Intriguingly, Metazoa had only sequences that belonged to Cluster 4 (of subfamily 1). Plants also displayed a great expansion in members of Cluster 4, but they also had small numbers of sequences from Cluster 1 and subfamily 2 (AzgA-like). Fungi had a rather balanced number of sequences from Cluster 1 and subfamily 2. Notably, Tobacco (*Nicotiana tabacum*) had the most (30) NAT/NCS2 proteins, the fungus *Basidiobolus meristosporus* had 16 members, whereas two lophotrochozoa (*Lingula unguis and Crassostrea gigas*) had 12 members each. Although it is conceivable that the percentages observed in this analysis may change depending on the species sampling, the general trends observed for each major taxonomic lineage are expected to hold.

## Conclusions

Based on a large-scale phylogenetic analysis of prokaryotic NAT/NCS2 proteins, a webserver has been developed that may scan whole proteomes and identify members of this family. The server classifies these members in various subfamilies and evolutionary clusters with certain substrate profiles and identifies conserved motifs that are related to function. An analysis of 9109 prokaryotic proteomes with our server revealed that the evolutionary lineages containing the largest numbers of NAT/NCS2 members are β- and γ-Proteobacteria, Bacilli, Clostridia, Actinobacteria and Fusobacteria. An analysis of 120 Eukaryotic proteomes also revealed that this server is fully capable of successfully analyzing this taxonomic lineage as well.

## Availability of supporting data

All supplementary data can be downloaded from the NAT/NCS2-hound server at: http://bioinf.bio.uth.gr/nat-ncs2/

## Author’s Contributions

AC and CN performed the phylogenetic analyses, PV developed the server, MB, VY, PL, ET gathered annotation and functional information. DM, SF and GDA conceived the study, supervised the students and prepared the manuscript.

## Funding

No funding was provided for the materialization of this work.

## Competing interests

None declared

